# PP2A-Cdc55 phosphatase coordinates actomyosin ring contraction and septum formation during cytokinesis

**DOI:** 10.1101/2020.11.27.400903

**Authors:** Yolanda Moyano-Rodríguez, David Vaquero, Odena Vilalta-Castany, Magdalena Foltman, Alberto Sanchez-Diaz, Ethel Queralt

## Abstract

Eukaryotic cells divide and separate all their components after chromosome segregation by a process called cytokinesis to complete cell division. Cytokinesis is highly regulated by recruitment of the components to the division site and through post-translational modifications such as phosphorylations. The budding yeast mitotic kinases Cdc28-Clb2, Cdc5, and Dbf2-Mob1 phosphorylate several cytokinetic proteins contributing to the regulation of cytokinesis. The PP2A-Cdc55 phosphatase regulates mitosis counteracting Cdk1- and Cdc5-dependent phosphorylation. This prompted us to propose that PP2A-Cdc55 could also be counteracting the mitotic kinases during cytokinesis. Here, we demonstrate that primary septum formation and actomyosin ring contraction are impaired in the absence of PP2A-Cdc55. In addition, by *in vivo* and *in vitro* assays, we show that PP2A-Cdc55 dephosphorylates the chitin synthase II (Chs2 in budding yeast) a component of the Ingression Progression Complexes (IPCs) involved in cytokinesis. Interestingly, the non-phosphorylable version of Chs2 rescues the asymmetric AMR contraction observed in *cdc55*Δ mutant cells. Therefore, timely dephosphorylation of the Chs2 by PP2A-Cdc55 is crucial for proper actomyosin ring contraction. These findings reveal a new mechanism of cytokinesis regulation by the PP2A-Cdc55 phosphatase and extend our knowledge in the involvement of multiple phosphatases during cytokinesis.

## Introduction

Cytokinesis is the final event of the cell cycle and mediates the physical separation of mother and daughter cells. It is a highly ordered and regulated process that is conserved among eukaryotes. Cytokinesis must be spatially and temporally coordinated with sister chromatids segregation to avoid problems in chromosome segregation –failures can lead to aneuploidies and are associated with tumorigenesis[1]. Detailed mechanistic studies are important to understand cytokinesis relevance in normal cellular functions as well as its impact on human diseases.

In fungi and animal cells, the cytokinetic machinery comprises two major elements: septin and contractile actomyosin rings. In *Saccharomyces cerevisiae*, the process takes place at the division site, a narrow region linking mother and daughter cells known as the bud neck. At the end of G1, the Cdc42 GTPase polarizes actin cables and patches towards the new bud site, and the septin ring is assembled[2,3].The septin ring is reorganized later in S phase and becomes an hourglass-like structure[4] that will split into two rings in anaphase[5–7]. Septins serve as an anchor for various cytokinesis-related proteins including the type II myosin heavy chain, Myo1[8]. Myo1, together with actin filaments and the essential and regulatory myosin light chains Mlc1 and Mlc2, shape the actomyosin ring (AMR)[5,8–10]. The AMR leads to the furrow ingression through its contraction and constriction[11,12].

Cytokinetic regulatory proteins such as Iqg1,Hof1, Inn1 and Cyk3 are recruited at the AMR[6,13–15], which mediate the activation of the chitin synthase II, Chs2[6,16–18]. Chs2 synthesizes the primary septum (PS), concomitantly with AMR constriction[19,20], and drives the second part of its contraction [21]. These proteins are part of the Ingression Progression Complexes (IPCs)[17,22], which coordinates primary septum formation, actomyosin ring contraction and ingression of the plasma membrane at the division site[16,17,23–26].

Once the PS has been synthesized, the two secondary septa (SS) will be formed by Chs3, glucan synthases (Fks1) and mannosyltransferases on both sides of the PS[20,27,28]. Finally, cytokinesis is completed when the daughter cell synthesizes hydrolases and chitinases to hydrolyze the remaining cell wall structures in between SS[29].

The timing of cytokinesis must be tightly regulated to ensure it only happens when anaphase is completed. Anaphase is regulated by two pathways: FEAR (CdcFourteen Early Anaphase Release) and MEN (Mitosis Exit Network). Both are coordinated to promote the activation of the Cdc14 phosphatase[30–33]. Cdc14 is responsible for Cdk1 (Cdc28 in budding yeast) inactivation and the dephosphorylation of Cdk1 targets during mitosis[34]. Several MEN kinases have been found to be involved in cytokinesis[35]. Polo-like kinase, Cdc5 regulates AMR formation and membrane ingression[36,37]. Dbf2, the downstream MEN kinase, phosphorylates Hof1 promoting its re-localization from the septin ring to the AMR[7,36]. Cdk1-Clb2 phosphorylates Iqg1 limiting the actin recruitment at the division site [38,39] and Chs2 promoting its retention at the endoplasmic reticulum (ER)[40–42]. Cdk1 also inhibits Iqg1 and Inn1 localization at the division site, and Inn1 interaction with IPCs[38,43,44]. Later on, MEN and Cdc14 phosphatase initiate cytokinesis by counteracting Cdk1 phosphorylation in budding yeast[39,41,44,45]. However, the inactivation of Cdk1-Clb2 activity during late anaphase is not sufficient to trigger cytokinesis[41,46], it is also necessary the Cdc14-dependent dephosphorylation of Iqg1, Inn1 and Chs2 for cytokinesis completion[39,41,44,45]. Chs2 translocates to the division site upon Chs2 dephosphorylation by Cdc14[41,47].

Recent work demonstrates that multiple phosphatases shape the phospho-proteome during mitotic exit[48,49], pointing out the contribution of other phosphatases such as PP2A-Rts1 and PP2A-Cdc55[48]. These observations suggest thatPP2A phosphatases could also be required for cytokinesis. Here, we propose that PP2A-Cdc55 regulates cytokinesis through the dephosphorylation of the IPC protein, Chs2. We have found that the PP2A-Cdc55 participates in the regulation of the phosphorylation state of Chs2 and dephosphorylates Chs2 *in vivo* and *in vitro*. In the absence of Cdc55, AMR contraction and the PS formation occur asymmetrically to one side of the bud neck supporting a role for PP2A-Cdc55 in cytokinesis regulation. Interestingly, the non-phosphorylable version of Chs2, *chs2-S133A* rescues the asymmetric AMR contraction in *cdc55Δ* mutant cells. The phospho-mimetic *chs2-S133E* promotes a longer Hof1 contraction time in wild-type cells, similarly to the absence of Cdc55, suggesting that hyperphosphorylation of Chs2 alters Hof1 residence time at the division site. In conclusion, PP2A-Cdc55 coordinates AMR contraction and PS formation via the dephosphorylation of Chs2 during cytokinesis.

## Results

### Defective septum formation and AMR contraction in the absence of Cdc55

It was described that PP2A-Cdc55 is localized at the division site during cytokinesis [50]. However, neither a role for PP2A-Cdc55 during cytokinesis was demonstrated nor the molecular mechanism by which PP2A-Cdc55 could regulate cytokinesis. This prompted us to study the cytokinesis phenotypes of the *cdc55Δ* mutant cells.

Septins act as scaffold platforms for many proteins and impose a diffusion barrier for regulating cell polarity, cell remodeling and cytokinesis [51]. Septins localization and organization are highly dynamic through the cell cycle [52,53]. We wondered whether septin structures were defective in the absence of Cdc55. Septins were visualized *in situ* by immunofluorescence staining of Cdc11 and Shs1-HA in asynchronous cells. The septin structures (septin rings, collar and double rings) were indistinguishable between *cdc55Δ* and wild-type cells (Fig. S1), suggesting that septin dynamics might not be greatly affected by the absence of Cdc55.

PP2A-Cdc55 regulates bud morphology through actin polarization and cell-wall synthesis [54]. Actin cytoskeleton polarization at the site of bud emergence is triggered at Start by the kinase activities of Cln1,2-Cdc28. Upon Clb2-Cdc28 kinase inactivation at the end of mitosis, the actin cytoskeleton is directed to the neck for the formation of the AMR [9,39]. We envisage the possibility that PP2A-Cdc55 influences also actin cytoskeleton during cytokinesis. To examine this possibility, actin filaments were stained with rhodamine-labelled phalloidin in cells progressing through mitosis and cytokinesis in *cdc55Δ* cells. Actin signal was found to be depolarized in metaphase-arrested cells and localized at the bud neck in anaphase-cytokinesis in wild-type and *cdc55Δ* cells (Fig. S2a). However, in a subpopulation of *cdc55Δ* cells, actin polarization at the new bud was observed before cytokinesis had been completed (Fig. S2b). We observed actin signals simultaneously at the division site during cytokinesis and at the new bud site in 12.5% of *cdc55Δ* mutant cells, whereas there was no premature actin re-polarization at the new bud site in wild-type cells (Fig. S2b). The premature actin localization indicates that actin re-polarization of the next cell cycle occurs before cell division in a fraction of *cdc55Δ* cells. Therefore, PP2A-Cdc55 could act to prevent actin re-polarization until cytokinesis is completed.

Cytokinesis in budding yeast is accomplished by the concerted action of the actomyosin contractile ring (AMR) and the septum formation. We investigated how these two processes occur in the absence of Cdc55. First, to investigate whether the actomyosin contractile ring is functional, we analyzed the localization of the Myo1-tdTomato fusion protein in cells progressing through mitosis and cytokinesis after synchronous release from the metaphase arrest by Cdc20 depletion. It has been described that Myo1 localizes to the division site immediately after budding. Accordingly, in metaphase-arrested cells Myo1-tdTomato was localized at the bud neck in control cells. Similarly, Myo1-tdTomato was also detected at the bud neck during metaphase in *cdc55Δ* mutant cells. In anaphase, the Myo1 signal is reduced, reflecting the contraction of the AMR, until the signal becomes a single dot and finally disappears (Fig. 1a). The dynamics of AMR contraction were similar in the two strains, but we observed that AMR contraction was asymmetric respect to the centripetal axis in 86% (N=69) of *cdc55Δ* cells (Fig. 1b). In order to check whether the asymmetric Myo1 signal is not due to an adaptive mechanism of the *cdc55Δ* deletion mutant, we investigated the Myo1 contraction after inducing the Cdc55 degradation during metaphase using an auxin-degradation system [55]. Upon Cdc55 degradation, Myo1 contraction was also mostly asymmetric (86% of cells) (Fig. 1c). The results indicate that the lack of PP2A-Cdc55activity promotes the asymmetric contraction of Myo1.

**Figure 1.**
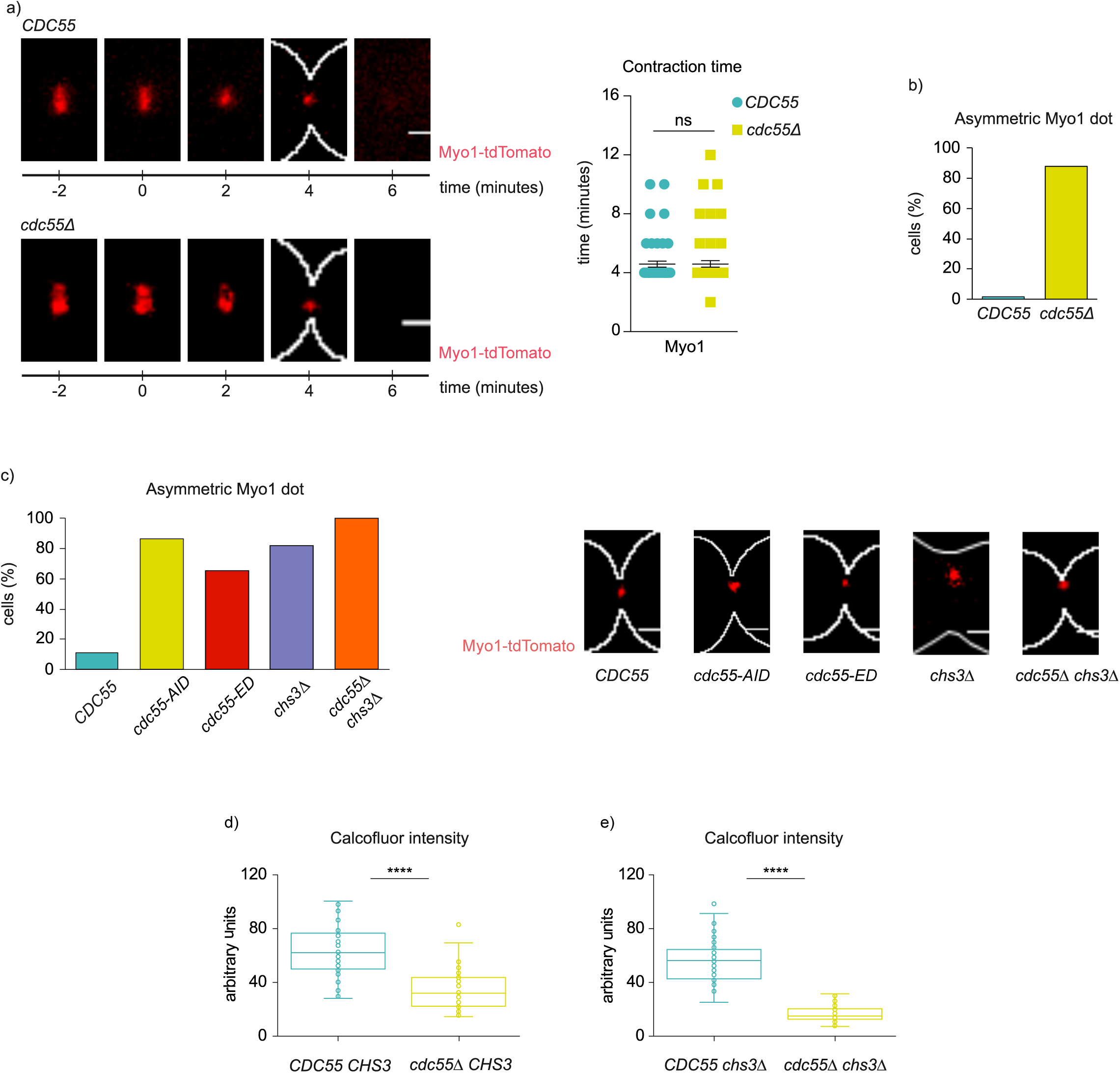
PP2A-Cdc55 regulates AMR constriction and septum formation. (a) Absence of Cdc55 promotes the asymmetry of Myo1upon AMR contraction. Strains Y1306 and Y1578 were arrested in metaphase and released into anaphase by Cdc20 depletion and re-addition, and time-lapse images were captured every 2 minutes. Spc42-GFP (spindle pole body protein) was used as the control for anaphase progression. Representative images of Myo1 in *CDC5*5 and *cdc55Δ* are shown (left panel). Quantification of the contraction time of Myo1 is depicted (right panel). (b) Quantification of the population of cells with asymmetric Myo1 constriction in *CDC55* (N=76) and *cdc55Δ* (N=69) from (a). (c) Myo1 asymmetric localization upon degradation or inactivation of Cdc55. Strains Y1652 (N=9), Y1653 (N=17), Y1788 (N=35), Y1605 (N=22) and Y1596 (N=24) were arrested in metaphase and released into anaphase by Cdc20 depletion and re-addition. *cdc55-aid* degradation was induced by the addition of 0.5 mM auxin for 3 hours during the metaphase arrest and auxin was maintained after Cdc20 re-addition. Time-lapse images were captured every 2 minutes. Quantification of the cell population with asymmetric Myo1 constriction (left panel), and maximum intensity z-projection images of Myo1 (right panel) are shown. Scale bar, 1 μm. (d-e) Chitin deposition at primary and secondary septa is regulated by PP2A-Cdc55. Strains Y1516 (N=46), Y1512 (N=49), Y1605 (N=82) and Y1596 (N=86) were arrested in metaphase by Cdc20 depletion and released into anaphase by Cdc20 re-induction. Calcofluor was added to visualize the septa upon the metaphase release. Student’s unpaired t-test was carried out using the Prism5 program.

We next examined whether Myo1 asymmetry could also be detected in the inactive version of Cdc55 (*cdc55-ED*). We synchronized cells at the metaphase-anaphase transition by Cdc20 depletion and analyzed the contraction of the Myo1-tdTomato. We observed that 65% of *cdc55-ED* mutant cells showed an asymmetric Myo1-tdTomato signal upon contraction (Fig. 1c). This result indicates that, similar to the absence of Cdc55, the non-functional Cdc55 results in the alteration of AMR contraction. Overall, we conclude that Myo1 recruitment and time of localization at the division site are not altered in the absence of Cdc55, although AMR contraction is defective in the absence of PP2A-Cdc55 activity. This asymmetric localization has been reported before in some IPCs mutants [26] and suggests a dysfunctional AMR. These results suggest that PP2A-Cdc55 is required for the correct function of AMR and for an efficient cytokinesis.

To assess whether the asymmetric AMR contraction phenotype was not affected by the synchronization method, we performed the assay synchronizing cells in G1 by alpha factor in absence of Cdc55. *cdc55Δ* cells enter mitosis with a delay due to compromised Cdk1 activity because of inhibitory Cdc28-Y19 phosphorylation [56]. To correct for this delay, we introduced the *cdc28_Y19F* allele, which is refractory to Cdk1 inhibition. The *cdc55Δ* cells containing *cdc28_Y19F* progressed normally through the cell cycle [57]. Again, we observed that Myo1-tdTomato constriction was asymmetric in 80% of the *cdc55Δ cdc28_Y19F* cell population (Fig. S3). Therefore, the asymmetric Myo1 signal in the absence of Cdc55 was observed independently of the synchronization method used.

To study septum formation, chitin deposition at the division site was analyzed by *in vivo* staining with calcofluor white and measured the fluorescence intensity of the incorporated calcofluor on living cells containing Myo1-tdTomato as a control for cytokinesis progression. We arrested cells at metaphase by Cdc20 depletion, released them into mitosis and took images 40-50 min after the release when we found cells at cytokinesis. Calcofluor intensity was then measured and quantified in wild-type and *cdc55Δ* cells (Fig. 1d). There was a 45% reduction in the intensity of the calcofluor staining in the absence of Cdc55 compared to control cells. Chitin is incorporated in primary septum (PS) and secondary septa (SS); therefore, we cannot distinguish at which septum the reduction occurs. For this reason, we repeated the calcofluor staining in cells containing a deletion for*CHS3*, the chitin synthase responsible for secondary septa formation [20]. As previously described, *chs3Δ* cells showed wider bud necks and cytokinetic defects provoking the formation of chains of cells [58]. A reduction of 70% in the calcofluor intensity was measured in the *chs3Δ* background in the absence of Cdc55 (Fig. 1e). Interestingly, we noticed that 82% (N=22) of the cells showed asymmetric Myo1 contraction in *chs3Δ* cells (Fig. 1c), indicating defects in cytokinesis [26]. The above results indicate that primary and secondary septa formation are reduced in cells lacking PP2A-Cdc55 activity, suggesting that PP2A-Cdc55 has a role coordinating AMR contraction with septum formation.

Primary septum formation and membrane invagination occur concomitantly. To determine whether ingression of the plasma membrane could occur in *cdc55Δ* cells, we performed time-lapse video microscopy of cells expressing the small G-protein Ras2 fused to 3 copies of GFP to study plasma membrane dynamics. The visualization of the plasma membrane at the site of division revealed no cytoplasm connection between mother and daughter cells after cytokinesis, confirming that cytoplasmic division is resolved in control and *cdc55Δ* cells (Fig. S4). In conclusion, *cdc55Δ* cells have an asymmetric AMR contraction and a reduction in septa formation, but nevertheless manage to complete cell division.

Next, to study the defects in septa formation we investigated the cytokinetic structure by transmission electron microscopy (TEM). We synchronized cells at the metaphase-anaphase transition by Cdc20 depletion and captured images 40-50 min after the synchronous release. Cells synthesizing the primary septum were identified and the structure of the septum was then examined. In wild-type cells, we observed PS formation from the cell wall following membrane invagination (Fig. 2a), as expected. Once the PS was finished, the two SS was formed on both sides of the PS (Fig. 2b). Finally, the PS is degraded, and the cells physically separated (Fig. 2c). Conversely, in *cdc55Δ* cells, the PS emerged from only one side of the division site (Fig. 2d). From the cells performing primary septum formation, we quantified 85% (N=25) with asymmetric PS in absence of Cdc55. Some cells showed aberrant, thicker structures with diverse morphologies that resemble the secondary septa-like structures, commonly referred as remedial septum (Fig. 2e). The remedial septum was first described in IPCs mutant cells, i.e., *myo1Δ*, *chs2Δ* and *chs3Δ* cells[24]. They are chitin structures reminiscent of SS that allow cells to complete cytokinesis when the PS is defective [24,59]. Strikingly, in *cdc55Δ* cells, the mother and daughter cells are finally able to separate physically, probably upon formation of the remedial-like septum. To further investigate this hypothesis, we repeated the experiments using a *chs3Δ* background in which no secondary septa is formed (Figs. 2g-i). Similar results were obtained, whence the *cdc55Δ chs3Δ* cells had asymmetric PS formation and a remedial-like septum was formed (Figs. 2j-l). These results suggest that remedial-like septum formation enables cytoplasm separation and cell division in the absence of Cdc55.

**Figure 2.**
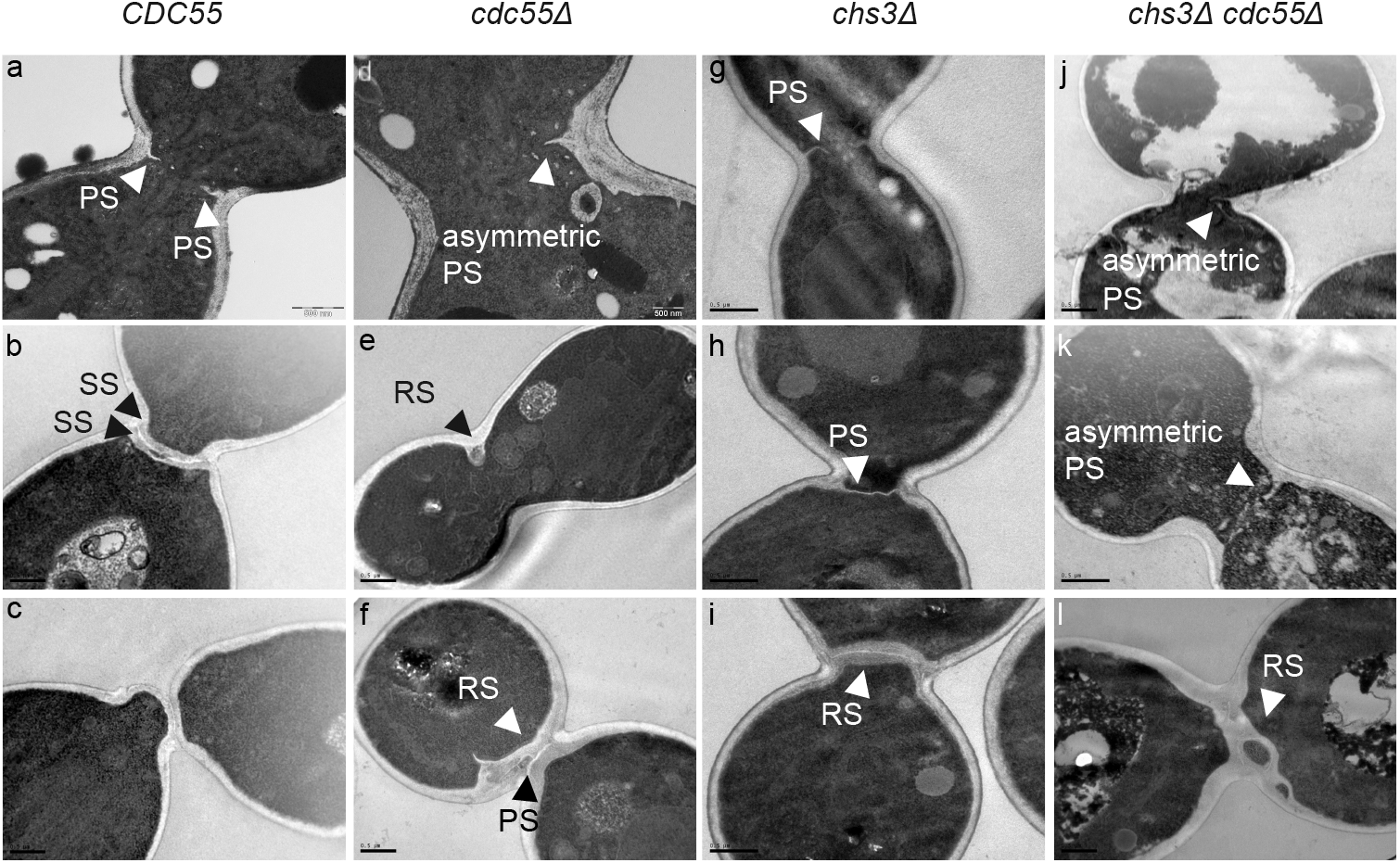
Asymmetric primary septum formation and appearance of the remedial-like septum in the absence of Cdc55. Strains Y1315, Y1318, Y1605, and Y1596 were arrested in metaphase and released into anaphase by Cdc20 depletion and re-addition. Representative images from TEM are shown. Scale bar, 0.5 μm. PS, SS and RS denote primary septum, secondary septa and remedial septum, respectively.

In summary, the asymmetric PS formation detected by EM and the asymmetric AMR contraction observed in *cdc55Δ* cells, indicates that PP2A-Cdc55 has a role regulating the correct coordination of the AMR contraction and PS formation.

### PP2A-Cdc55 regulates the dephosphorylation of Chs2

The Ingression Progression Complexes (IPCs) coordinate AMR constriction, plasma membrane ingression and septum formation[17,18]. We envisaged that PP2A-Cdc55 may have a role during cytokinesis through the dephosphorylation of IPCs proteins[60]. We initially investigated the genetic interactions between *cdc55Δ* and IPCs mutants. We prepared double mutants with *cdc55Δ* and degron-conditional mutants to induce protein degradation for the IPCs subunits[17,61]. No differences in cell growth were observed on control plates (Fig. 3a). However, the viability of the double mutants *cdc55Δ hof1-aid* and *cdc55Δ td-cyk3-aid* was impaired under restrictive conditions (presence of auxin) (Fig. 3a). The synthetic sick interactions found between *cdc55Δ* and *hof1-aid* or *td-cyk3-aid* degron mutants indicate that Cdc55 is functionally related to the IPC subunits, suggesting that Cdc55 is involved in the regulation of cytokinesis.

**Figure 3.**
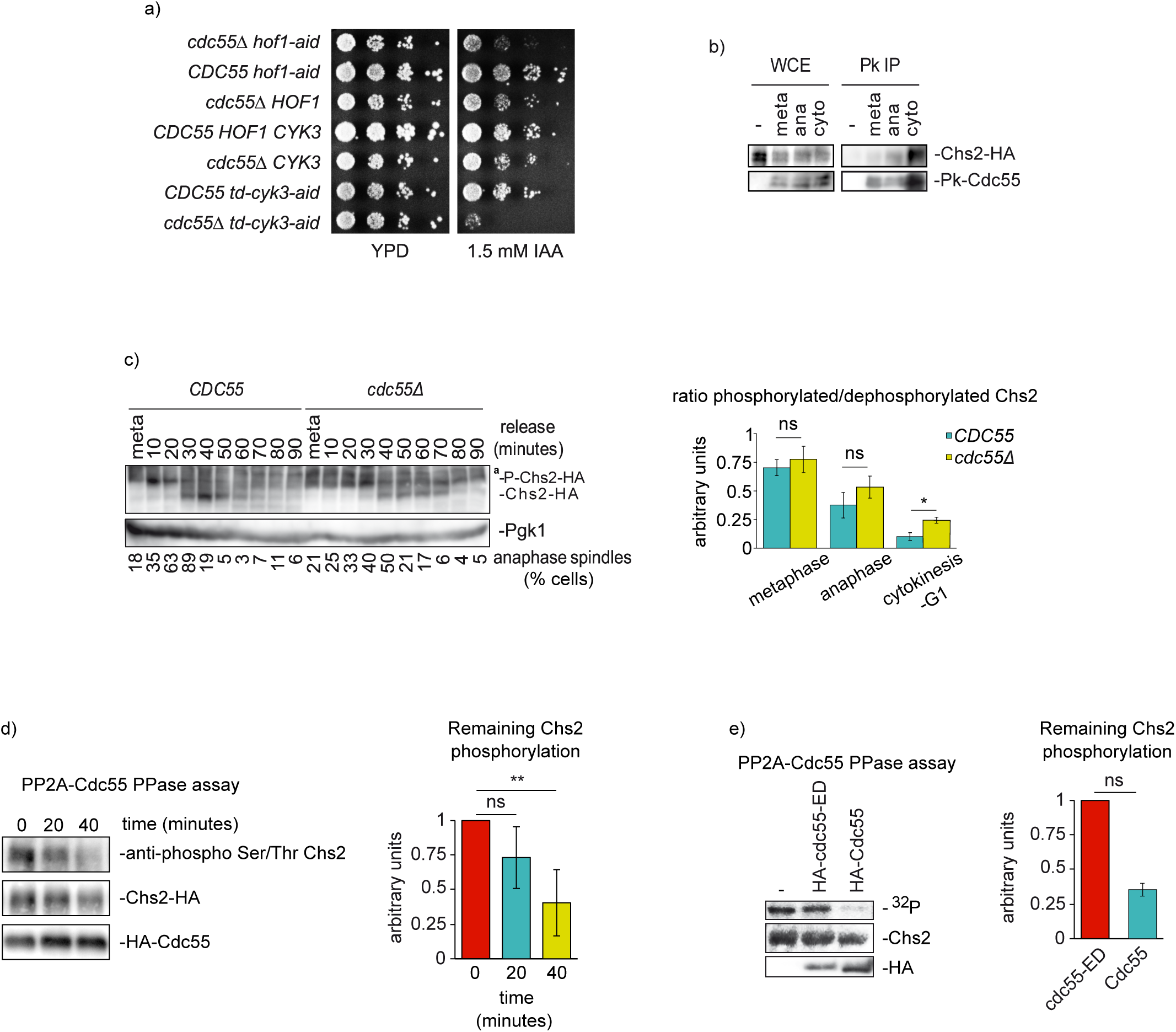
PP2A-Cdc55 dephosphorylates Chs2 during cytokinesis. (a) The double mutants *cdc55 hof1* and *cdc55cyk3* are synthetically sick. Serial dilutions ofW303, Y844, Y1728, Y1730, Y1747 and Y1749 strains were spotted on YPD plates with and without 1.5 mM auxin (IAA). Cells were grown at 25°C for 2-3 days. (b) PP2A-Cdc55 forms a complex with Chs2. Strain Y1567 was synchronized into anaphase progression by Cdc20 depletion and re-addition. Strain Y1318 without the Pk-tag was used as negative control. Protein extracts were prepared at metaphase, anaphase, and cytokinesis and Cdc55 was immunoprecipitated using the anti-Pk antibody. Co-purification of Chs2 was analyzed by western blot. (c) PP2A-Cdc55 regulates Chs2 dephosphorylation. Strains Y1318 and Y1315 were arrested in metaphase and released into anaphase by Cdc20 depletion and re-addition. Chs2 phosphorylation was analyzed by western blot in Phos-tag gels. Unspecific bands were detected for Chs2 and marked as (^a^). Pgk1 levels were used as a loading control. Mitosis progression was followed by analyzing the anaphase spindle elongation by *in situ* immunofluorescence. At least 100 cells were scored at each time-point. Quantification of the western blots was performed using Fiji Software and means and SEMs are represented. Student’s unpaired t-test analyses were carried out using the Prism5 program. (d) Chs2 is dephosphorylated by PP2A-Cdc55. Metaphase arrested cells of the strain Y695 were used to purify the PP2A-Cdc55 complex by TAP purification. Chs2-HA was purified from metaphase arrested cells of strain Y1318. Purified Chs2-HA was incubated with the PP2A-Cdc55 complex at the indicated times and the Chs2 phosphorylation levels were detected by western blot using the anti-phospho Ser/Thr antibody. Representative images of one phosphatase assay are shown. Protein levels were quantified using Fiji software. Quantifications of the remaining Chs2 phosphorylation signal normalized to the amount of Cdc55 and Chs2 are shown. Means and SEMs of three phosphatase assays are represented. Student’s unpaired t-test analysis was carried out using the Prism5 program. (e) PP2A-Cdc55 dephosphorylates Chs2 *in vitro*. Strains Y824, Y1652 and Y1653 were arrested in metaphase by Cdc20 depletion. Cdc55 and *cdc55-ED* were purified by immunoprecipitation with HA antibody. Strep-tag-Chs2-1-629 purified from *E. coli* and previously phosphorylated by Cdk1-Clb2 in a radioactive kinase assay was used as substrate. Representative images of the phosphatase assays are shown. Radioactive signals were detected using a multi-purpose imaging plate in a Typhoon FLA950 apparatus (GE healthcare). Protein levels were quantified using Fiji software. Quantifications of the remaining phosphorylation signals normalized with respect to the amount of immunopurified HA-Cdc55 or HA-*cdc55-ED* are shown. Means and SEMs of three phosphatase assays are represented. Student’s unpaired t-test analysis was carried out using the Prism5 program.

To determine whether the IPCs proteins physically interact with PP2A-Cdc55, we performed co-immunoprecipitation experiments. We synchronized cells in metaphase as described above, and co-purification of Chs2 during anaphase and cytokinesis were detected in Cdc55 immunoprecipitates (Fig. 3b). Taken together, these results suggest that Chs2 forms a complex with PP2A-Cdc55 during progression through mitosis and cytokinesis.

To study the functional link between Cdc55 and the IPCs, we studied whether PP2A-Cdc55 regulates the dephosphorylation of IPC proteins. We analyzed the phosphorylation status of the IPC proteins in the absence of Cdc55 (Fig 3c and S5a-d) and observed an increase of Chs2 phosphorylation levels during cytokinesis. First, we synchronized cells at metaphase and released them synchronously into anaphase to visualize the phosphorylation of Chs2 during mitosis and cytokinesis. Mitotic spindle was stained by tubulin immunofluorescence and used as marker of cell-cycle progression. Cdc14 release from the nucleolus was also determined as control of mitosis progression. Cdc14 was prematurely release in metaphase in absence of Cdc55 as previously published[57]. During the time-course, Chs2 protein presented slow migration isoforms at metaphase and early anaphase in control cells (Fig. 3c meta-20 min). From anaphase until early G1 (Fig. 3c; 30-50 min), two Chs2 isoforms were detected. Since Chs2 presented many different migrating bands, we performed an alkaline phosphatase experiment to check which corresponds to phosphorylation. In presence of alkaline phosphatase, the slower Chs2 migrating bands collapse into the faster migrating band indicating that the upper bands are phosphorylation events (Fig. S5e). Remarkably, in *cdc55Δ* mutant cells the slower-migrating isoforms are maintained throughout the time-course, indicating that Chs2 was not efficiently dephosphorylated during late anaphase/cytokinesis (Fig. 3c; 40-70min). These results suggest that Chs2 phosphorylation levels are altered in the absence of Cdc55 during cytokinesis.

Later, to explicitly test whether Chs2 is dephosphorylated by PP2A-Cdc55, we measured PP2A-Cdc55 phosphatase activity against Chs2 after PP2A-Cdc55 purification from metaphase-arrested cells when PP2A-Cdc55 is active[57]. Immunopurified-Chs2 was incubated with purified PP2A-Cdc55 and Chs2 dephosphorylation was visualized using an anti-phospho Ser/Thr antibody (Fig. 3d). A 60% reduction in the Chs2 phosphorylation signal was measured after 40 min of incubation in presence of PP2A-Cdc55. As negative control, the phosphatase assay was done in the same conditions using separase (Esp1) as substrate. Immunopurified-Esp1 was incubated with purified PP2A-Cdc55 and the phosphorylation levels of Esp1 visualized with the anti-phospho Ser/Thr antibody (Fig. S5f). Esp1 phosphorylation signal was maintained during the experiment indicating that PP2A-Cdc55 is not able to dephosphorylate Esp1 in the same conditions used for Chs2. This result suggests that PP2A-Cdc55 dephosphorylation occurs specifically over Chs2.

Finally, we repeated the PP2A-Cdc55 phosphatase assay using a recombinant fragment of Chs2 (1-629) purified from *E. coli* and previously phosphorylated by Cdk1-Clb2. *In vitro* ^32^P-phosphorylated Chs2 substrate was incubated with Cdc55 immunoprecipitated from control cells expressing Cdc55 and *cdc55-ED* inactive version of PP2A-Cdc55[62]. A reduction of 84% in the Chs2 phosphorylation signal was detected in the presence of control Cdc55. The *cdc55-ED* mutant version was not able to dephosphorylate Chs2 significantly (Fig. 3e). These results strongly suggest that Chs2 is dephosphorylated by PP2A-Cdc55 *in vivo* and *in vitro*.

### PP2A-Cdc55 regulates IPCs symmetric localization and residence time at the division site

IPCs proteins form a complex whose function is to coordinate AMR contraction, plasma membrane ingression and PS formation[17,18]. For this reason, we examined whether the absence of Cdc55 disturbs the IPCs subunits proper or timely localization at the division site. To do this, we studied the localization of the IPCs subunits by time-lapse microscopy in the absence of Cdc55. We arrested cells in metaphase by Cdc20 depletion and captured images every 2 minutes after synchronous release into anaphase by Cdc20 re-induction. Iqg1, Inn1, Hof1, Cyk3 and Chs2 were contracted asymmetrically at the division site in *cdc55Δ* cells (Fig. 4a), consistent with the asymmetry of Myo1 described above. These findings indicate that the lack of PP2A-Cdc55 activity provokes a defective contraction of the AMR, resulting in defective PS formation.

**Figure 4.**
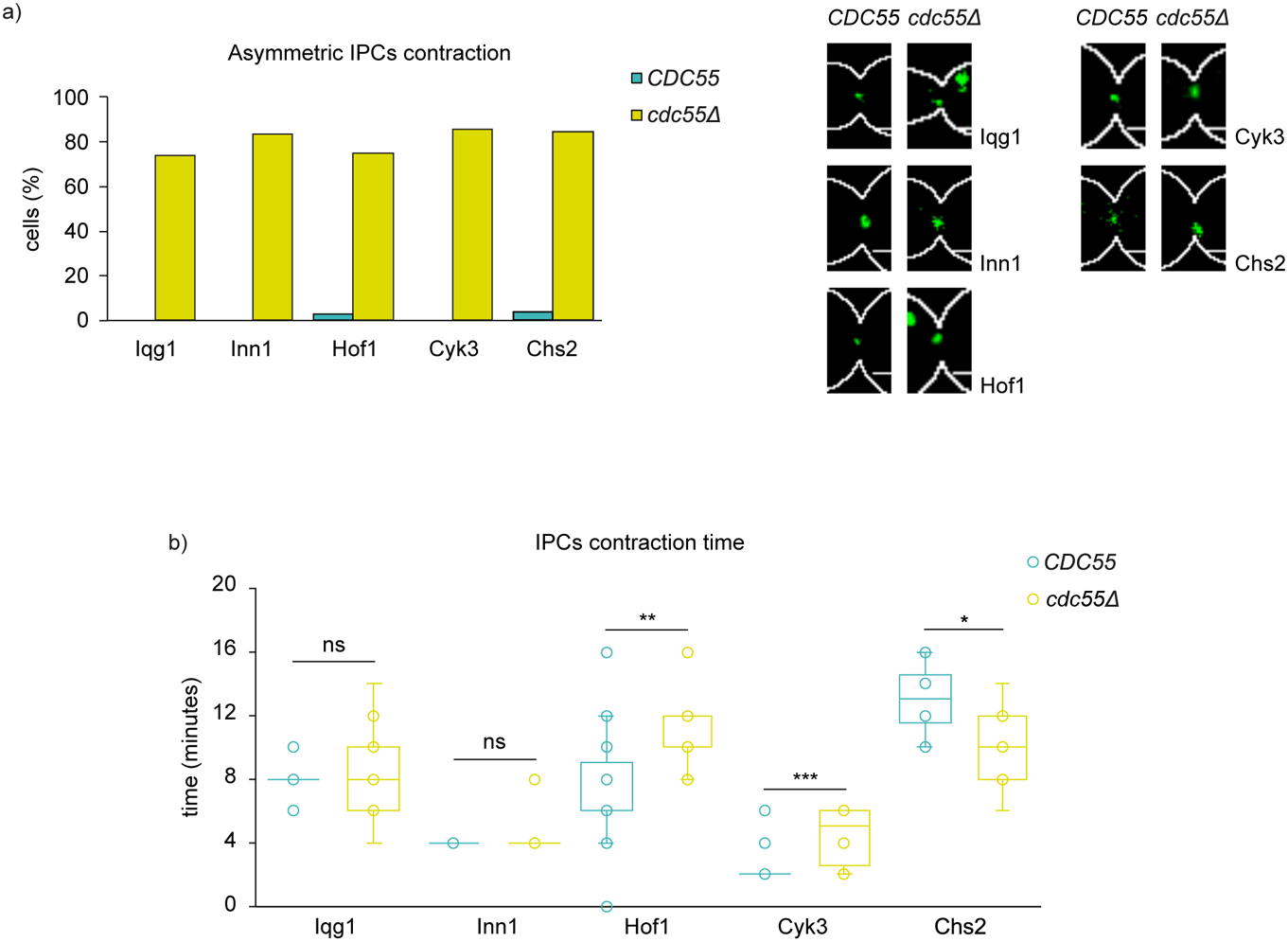
PP2A-Cdc55 is required for proper IPCs localization and contraction at the division site. Strains Y1572 (N=27), Y1606 (N=28), Y1454 (N=14), Y1608 (N=41), Y1306 (N=32), Y1578 (N=12), Y1574 (N=33), Y1604 (N=11), Y1576 (N=10) and Y1575 (N=8) were synchronized in mitosis by Cdc20 depletion and re-addition, and time-lapse images were captured every 2 minutes. Spc42-GFP and Myo1-tdTomato were used as a control for cytokinesis progression. (a) Percentages of cells with IPCs asymmetric Myo1 contraction are shown (left panel). Representative images of GFP-tagged proteins in *CDC5*5 and *cdc55Δ* are depicted (right panel). (b) Quantifications of the IPCs contraction times. Student’s unpaired t-tests were performed using the Prism5 program. Scale bar, 1 μm.

We then investigated the contraction and residence time of the IPCs subunits (Fig. 4b). No differences were detected in the residence time at the bud neck in Iqg1-GFP and Inn1-GFP proteins (Fig. 4b and Fig. S6a-b). However, we observed that Hof1-GFP and Cyk3-GFP signals took longer to complete contraction and to disappear in *cdc55Δ* cells (Fig. 4b and Fig. S6c-d), further indicating that cytokinesis is affected in absence of PP2A-Cdc55. The longer localization of Hof1 and Cyk3 at the division site could be a result of the impaired AMR contraction.

On the contrary, the Chs2-GFP residence time in *cdc55Δ* mutant cells was reduced compared to wild-type cells (Fig. 4b and S6e). Therefore, this result suggests that PP2A-Cdc55 regulates Chs2 localization dynamics, probably through the dephosphorylation of Chs2.

### Timely Chs2 dephosphorylation by PP2A-Cdc55 is required for proper AMR contraction

Our results suggest that PP2A-Cdc55 regulates Chs2-dependent processes during cytokinesis, such as PS formation and AMR contraction, and that participates in the dephosphorylation of Chs2. Therefore, PP2A-Cdc55 might contribute to cytokinesis regulation by controlling the phosphorylation levels of Chs2.

To screen for proteins regulated by the PP2A-Cdc55 phosphatase we studied the phosphoproteome in absence of Cdc55 by a quantitative analysis based on SILAC labeling. Phosphopeptides were enriched by Tish enrichment[63] and identified by LC-MS/MS. The screening revealed that a phosphopeptide corresponding to Chs2 protein was hyperphosphorylated in *cdc55Δ* mutant cells. The Chs2 peptide contained one Cdk1 minimal S/TP site: S133 (Fig. 5a). The phosphosite was detected with the highest confidence (pRS site probability 99.9% and p-value < 0.00001) and was also identified in a second phospho-proteomic study for *cdc55Δ* mutant cells[49]. The S133 is one of the 6 Cdk1 consensus sites (S/T-P) previously reported to be phosphorylated by Cdk1[64]. Chs2’s residues S14, S60, S69 and S100 containing the SP consensus sequence were described to be efficiently dephosphorylated by Cdc14,and the phospho mimetic mutant at those four sites, *chs2-S4E*, promotes Chs2 constitutive localization in the ER [41,47]. Cdc14 is active as a phosphatase and prematurely released from the nucleolus in *cdc55Δ* mutant cells[57,65]; therefore, the S133 hyperphosphorylation is a consequence of the lack of PP2A-Cdc55 activity since Cdc14 is not defective in absence of Cdc55.

**Figure 5.**
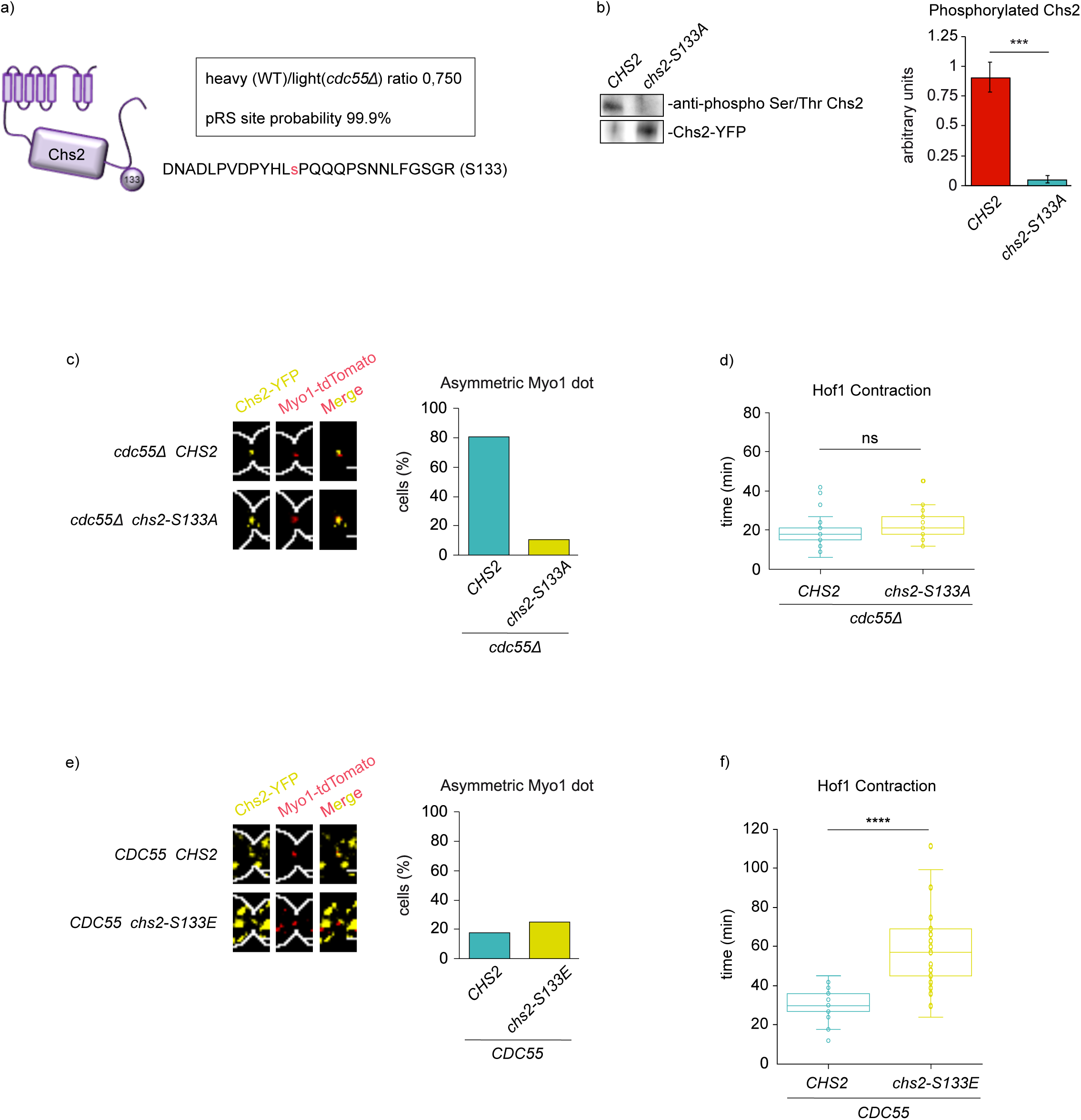
Chs2 phospho-mutants at S133 regulate AMR contraction and Hof1 residence time at the division site. (a) Hyperphosphorylation of Chs2 in S133 was detected by mass spectrometry in absence of Cdc55. Representation of the Chs2 peptide detected by mass spectrometry containing the S133. (b) Chs2 phosphorylation is greatly reduced in the *chs2-S133A* mutant. Strains Y1901 and Y1900 were arrested in metaphase and *GAL1-CHS2* versions were induced by galactose addition for 3 h. Protein extracts were prepared and Chs2 and *chs2-S133A* purified using GFP-trap columns. Chs2 phosphorylation levels were detected by western blot using the anti-phospho Ser/Thr antibody. Representative blots of one assay are shown (left panel). Western blot signals were quantified using Fiji software. Quantifications of the Chs2 phosphorylation levels normalized to the amount of Chs2 from three independent experiments are shown (right panel). Means and SEMs are represented. Student’s unpaired t-test analysis was performed using the Prism5 program. (c) The non-phosphorylable *chs2-S133A* version rescues the asymmetric AMR contraction in the *cdc55Δ* mutant cells. Strains Y1896 (N=25) and Y1898 (N=31) were arrested at metaphase by Cdc20 depletion. *GAL1-CHS2* versions were induced by galactose addition 3 hours before synchronous release into anaphase by Cdc20 re-introduction, and time-lapse images were captured every 2 minutes to visualize Myo1 and Chs2. Representative images of Myo1 and Chs2 are shown (left panel). Percentages of cells with asymmetric Myo1 are represented (right panel). (d) Similar Hof1 contraction times of Chs2 and *chs2-133A* in *cdc55Δ* mutant cells. Strains Y1912 (N=36) and Y1913 (N=38) were arrested at metaphase by Cdc20 depletion and *GAL1-CHS2* versions induced as in (c). Time-lapse images were captured every 3 minutes to visualize Hof1 and Chs2. Quantifications of the Hof1 contraction times are represented. Student’s unpaired t-tests were performed using the Prism5 program. (e) The phospho-mimetic *chs2-S133E* is not able to induce asymmetric Myo1 signal. Strains Y1895 (N=35) and Y1893 (N=29) were arrested at metaphase by Cdc20 depletion, *GAL1-CHS2* versions induced as in (c), and time-lapse images were captured every 2 minutes to visualize Myo1 and Chs2. Representative images of Myo1 and Chs2 are shown (left panel). Percentages of cells with asymmetric Myo1 are represented (right panel). (f) The phospho-mimetic *chs2-S133E* version promotes longer Hof1 residence time at the division site. Strains Y1909 (N=25) and Y1911 (N=26), were arrested at metaphase by Cdc20 depletion and *GAL1-CHS2* versions induced as in (c). Time-lapse images were captured every 3 minutes to visualize Hof1 and Chs2. Quantifications of the Hof1 contraction times are represented. Student’s unpaired t-tests were performed using the Prism5 program.

To investigate whether the S133 is a critical Chs2 phosphorylation site, we introduced a non-phosphorylable version of Chs2, *GAL1-chs2-S133A-YFP*, in wild-type cells and analyzed the phosphorylation levels in Chs2-YFP immunoprecipitates using the anti-phospho Ser/Thr antibody. Chs2 phosphorylation was detected in control cells expressing the wild-type version of Chs2-YFP (Fig. 5b). By contrast, the signal was greatly reduced in the immunoprecipitation of the non-phosphorylable version *chs2-S133A* (Fig. 5b). This result suggests that the S133 of Chs2 is phosphorylated.

If the cytokinetic defects observed in *cdc55Δ* mutant cells are consequence of the hyperphosphorylation of Chs2, the non-phosphorylable Chs2 version should rescue the asymmetric Myo1 contraction. For this reason, we introduced the non-phosphorylable version *GAL1-chs2-S133A-YFP* in wild-type and *cdc55Δ* mutant strains. Cells were synchronized by Cdc20 depletion and re-addition and *GAL1-CHS2* versions were induced prior to metaphase release. We followed Myo1-tdTomato and Chs2-YFP dynamics during mitosis and cytokinesis by time-lapse microscopy. Myo1 signal was asymmetric in the presence of the control Chs2-YFP (80% of cells; N=25), as expected in a *cdc55Δ* background (Fig. 5c). Remarkably, in the *chs2-S133A* non-phosphorylable version Myo1 became symmetric in 90% of the cells(N=31) (Fig. 5c). This indicates that the non-phosphorylable Chs2 mutant rescued the asymmetric localization of Myo1 in absence of Cdc55, Next, we wondered whether the *chs2-S133A* version also rescued the longer Hof1 contraction time observed in *cdc55Δ* cells (see Fig. 4b) by investigating the Hof1-mCherry dynamics. Interestingly, Hof1 contraction was similar in Chs2 and *chs2-S133A* in absence of Cdc55 (Fig. 5d). In conclusion, in absence of Chs2 phosphorylation at S133A, Myo1 signal is symmetric and Hof1 residence time is normal, suggesting that the S133 is the main dephosphorylated residue regulated by PP2A-Cdc55.

Finally, to investigate whether the hyperphosphorylation of Chs2 is sufficient to alter AMR symmetric contraction we introduced the phospho-mimetic *chs2-S133E* version in wild-type cells (Fig. 5e). No significant increase in the asymmetric Myo1 signal in the presence of *chs2-S133E* was observed suggesting that Chs2 phosphorylation is not sufficient to promote asymmetric AMR contraction. Nevertheless, an increase in the Hof1 residence time was observed in the phospho-mimetic *chs2-S133E* version (Fig. 5f), similarly to what we observed in *cdc55Δ* mutant cells (compare to Fig. 4b). These results suggest that PP2A-Cdc55 has a role in cytokinesis regulating Chs2 phosphorylation.

Overall, our findings demonstrate that the lack of PP2A-Cdc55 activity provokes an asymmetric contraction of the AMR and a defective PS formation that depends on the tight regulation of Chs2 phosphorylation. Therefore, Chs2 dephosphorylation by PP2A-Cdc55 is essential for maintaining proper AMR contraction and septum formation; both required for an efficient cytokinesis.

## Discussion

### PP2A-Cdc55 regulates Chs2 phosphorylation and its localization dynamics at the division site

The localization and function of the IPCs subunits(Iqg1, Myo1, Hof1, Cyk3, Inn1 and Chs2 proteins), which coordinate AMR contraction, plasma membrane ingression and PS formation, are tightly regulated by phosphorylation[43,44,66]. To date, only two phosphatases, Cdc14 and PP2A-Rts1, are known to have a role in cytokinesis[67], and it has been described that Cdc14 dephosphorylates several subunits of the IPCs proteins such as Chs2, Inn1 and Hof1[39,41,44,46]. A putative role for PP2A-Cdc55 in cytokinesis was also proposed because of its elongated morphology and multinucleated cells at low temperatures in *cdc55Δ* cells[68]. In addition, PP2A-Cdc55 is localized at the division site during cytokinesis[50]. However, the role of PP2A-Cdc55 during cytokinesis was not demonstrated and the molecular mechanism by which PP2A-Cdc55 regulates cytokinesis is still unknown.

Here, we describe a new function of PP2A-Cdc55 phosphatase in the regulation of cytokinesis. In summary, PP2A-Cdc55 regulates the dephosphorylation of Chs2 and it is involved in proper AMR contraction and septum formation. PP2A-Cdc55 dephosphorylates Chs2 *in vivo* and *in vitro*, indicating that it is a PP2A-Cdc55 target. The lack of dephosphorylation of Chs2 in absence of Cdc55 might be responsible of the defects seen in IPCs (Hof1) residence time (Fig. 4b) and AMR asymmetric contraction, since the *chs2-S133A* (Fig. 5) non-phosphorylable mutant rescue the asymmetric Myo1 signal in *cdc55Δ* mutant cells. In contrast, the phospho-mimetic mutant *chs2-S133E* is not sufficient to induce an asymmetric Myo1 signal in control cells, although provokes an increase in Hof1 residence time at the division site.

These results suggest that timely dephosphorylation of Chs2 by PP2A-Cdc55 is crucial for proper AMR contraction. Since the IPCs coordinate AMR constriction, plasma membrane ingression and septum formation, we propose that PP2A-Cdc55 is also involved in this efficient mechanism of coordination through Chs2 dephosphorylation (Fig. 6).

**Figure 6.**
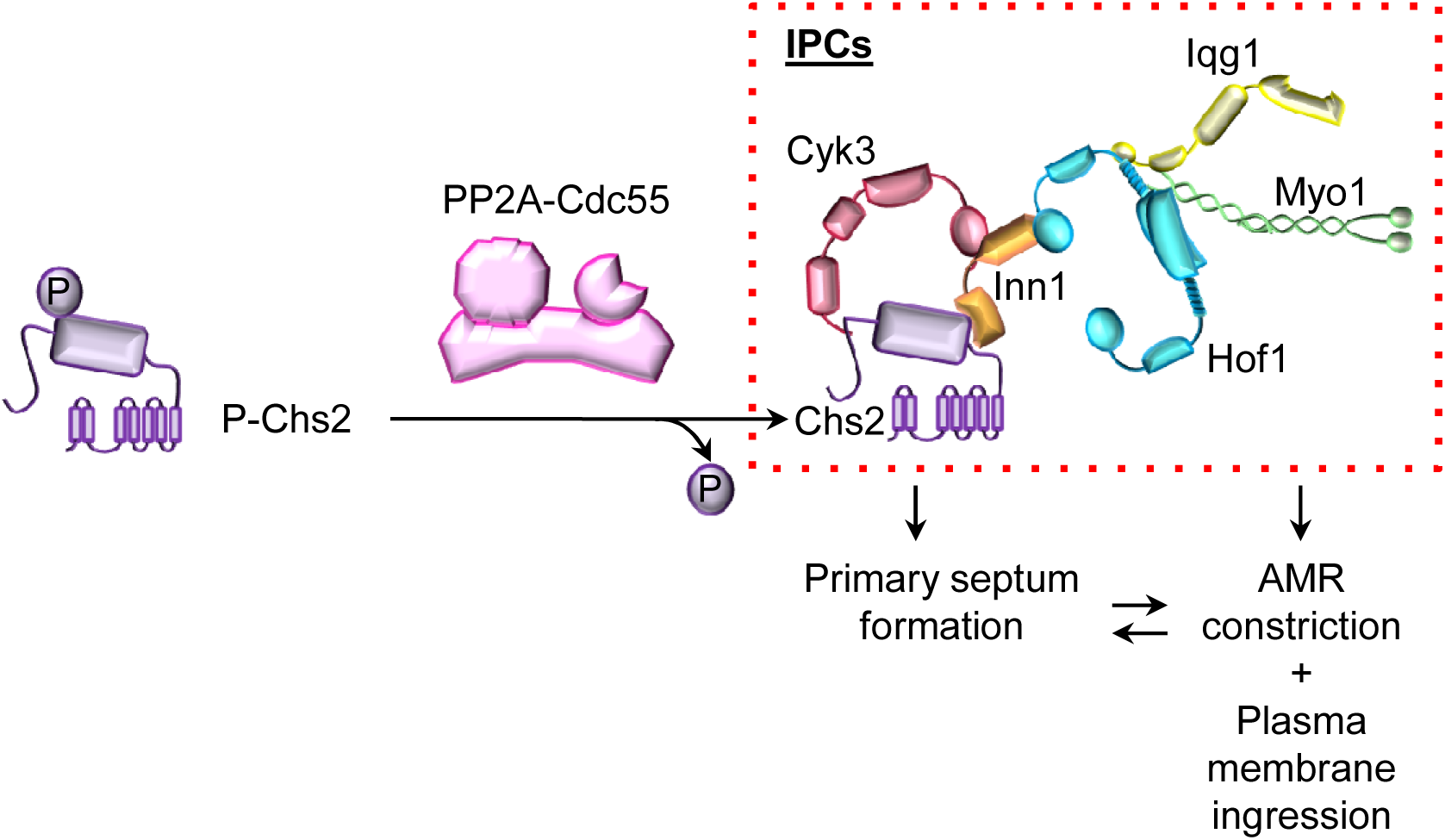
Model for PP2A-Cdc55 regulation of actomyosin ring (AMR) contraction, primary septum (PS) and secondary septa (SS) formation. Dephosphorylation of Chs2 by PP2A-Cdc55 coordinates AMR contraction and primary septum formation resulting in an increased Hof1 residence time at the division site.

### The role of PP2A-Cdc55 during cytokinesis

In the absence of Cdc55, Myo1 was constricted and unanchored in a timely fashion, but the constriction was displaced from the central axis, becoming asymmetric (Fig. 1a). This phenotype is characteristic of IPCs mutants[6,26,52] and denotes a dysfunctional AMR.

On the other hand,Hof1 asymmetric PS formation was described in *chs2Δ* cells[6]; but not *vice versa*[6,17], suggesting that Hof1 also influences PS formation. Therefore, defective PS formation in absence of Cdc55 could also contribute to the asymmetry observed in the other IPCs proteins.

In addition, PP2A-Cdc55 regulates septum formation. We observed that chitin incorporation was severely reduced in *cdc55Δ* and *cdc55Δ chs3Δ* cells (Fig. 1d), indicating that Chs2 activity might be impaired. Interestingly, the electron microscopy study demonstrated that the PS appears only at one side of the division site during its formation in absence of Cdc55 (Figs. 2d and j). Later, a remedial-like septum that contains even cytosol fractions embedded in the septa is detected in absence of Cdc55 (Fig. 2f and l). The remedial septum formation was previously described in IPCs and MEN mutants with defective PS formation[13,18,69,70]. Chs3 glucan synthases and mannosyltransferases are responsible for the SS formation and the construction of the remedial septum[27,28,59]. However, the *chs3Δ* mutant is still viable and capable of divide, suggesting that additional chitin synthases might be able to incorporate chitin in the absence of Chs3. This may explain why the remedial-like septum is still formed in the *cdc55Δ chs3Δ* double mutant (Fig. 2i and l).

Recently, it has been shown that multiple phosphatases shape the phospho-proteome during mitotic exit[48,49,71]. Notably, Hof1 and Chs2 phosphosites have been identified in *cdc55* and *cdc14* mutants[48]. We demonstrated that PP2A-Cdc55 regulates cytokinesis through the Chs2 dephosphorylation and it was reported that Cdc14-dependent dephosphorylation is required for Chs2 release from the ER[41], supporting different and specific contribution to cytokinesis of several phosphatases. We observed hyperphosphorylations of Hof1 and Inn1 during cytokinesis in absence of Cdc55 (Fig. S5a-b), suggesting that PP2A-Cdc55 might also regulates other IPCs proteins. Further investigations will be required to understand the coordinated regulation of multiple phosphatases during the cell cycle.

## Materials and methods

### Yeast strains, plasmids and cell-cycle synchronization procedures

All yeast strains used in this study were derivatives of W303 and are listed in Table S1. Epitope tagging of endogenous genes and gene deletions were performed by gene targeting using polymerase chain reaction (PCR) products[72]. Cell synchronization using α-factor and metaphase arrest by Cdc20 depletion and entry into synchronous anaphase by Cdc20 re-introduction were performed as previously described[73]. To obtain the *pGAL1-CDC55* construct, the DNA fragment containing the *GAL1* promoter was cut with SpeI and subcloned into the *CDC55* containing plasmid previously digested with NheI.

All the *cdc55Δ* mutant strains were freshly prepared by transformation or by crossing strains to avoid accumulation of suppressor mutations. The introduction of the *cdc55Δ* deletion was determined by PCR-genotyping and observation of the slow growth and elongated morphology under the microscope.

### Recombinant protein purification

All plasmids were freshly introduced into BL21 *E. coli*. Cells were grown in LB medium and protein expression was induced with 0.1 mM isopropyl β-D-1-thiogalactopyranoside (IPTG) at 25°C overnight. Collected cells were washed with PBS1 and frozen for at least 30 minutes.

Cells containing His_6_-Cyk3, His_6_-Inn1 or His_6_-Hof1 plasmids[17] were resuspended in cold lysis buffer (30 mM Tris-HCl, pH 8, 300 mM NaCl, 30 mM imidazole, 0.1% NP40, 10 mM β-mercaptoethanol, 1 mM PMSF, complete EDTA-free tablet (Roche)) and sonicated for 6 cycles of 1 min at 25 µm amplitude. Protein extracts were clarified and incubated with Ni-NTA magnetic beads (Thermo Fisher) at 4°C for 1 hour. Beads were washed with 10 volumes of lysis buffer and protein eluted in PBS, 5 mM EDTA, 5 mM DTT, 0.1% NP40, 500 mM imidazole at 4°C for 30-60 minutes.

Cells containing StreptagIII-Chs2_1-629 plasmid[17] were resuspended in cold lysis buffer (50 mM Tris-HCl, pH 8.0, 10% glycerol, 0.1% NP-40, 10 mM MgCl_2_, 300 mM NaCl, 5 mM β-mercaptoethanol, 10% BugBuster® (Millipore), 1 mM PMSF, complete EDTA-free tablet (Roche)) containing 5 U/mL of nuclease (Pierce), incubated in a rotatory wheel at 25°C for 20 minutes. Protein extracts were clarified and incubated with Strep-tactin® Superflow® resin (Iba Life science) at 4°C for 1 hour. Beads were washed with ten volumes of lysis buffer (without BugBuster®), and protein was eluted in 50 mM Tris-HCl, pH 8.0, 10% glycerol, 0.1% NP-40, 10 mM MgCl_2_, 150 mM NaCl, 5 mM β-mercaptoethanol, 2.5 mM desthiobiotin at 4°C for 1 hour.

### Co-immunoprecipitation, kinase assays and phosphatase assays

Co-immunoprecipitation assays were performed using 10^8^yeast cells, which were resuspended in lysis buffer (50 mM HEPES-KOH pH 7.5, 70 mM KOAc, 5 mM Mg(OAc)_2_, 10% glycerol, 0.1% Triton X-100, 8 μg/mL of protease inhibitors (leupeptin, pepstatin, aprotinin), 1 mM PMSF, 1.25mg/ml of benzamidin, 1 tablet of complete protease inhibitor without EDTA (Roche), 4 mM of phosphatase inhibitors (β-glycerophosphate, NaF and NEM) and 1 tablet of PhosStop (Roche)). Lysates were obtained by mechanical lysis using glass beads in a Bertin disrupter (6 cycles of 10 s at 5,000 rpm). Protein extracts were clarified by centrifugation and incubated with α-Pk clone SV5-Pk1 (AbSerotec) orα-HA clone 12CA5 (Roche) antibodies for 1 hour. Protein extracts were then incubated for 1 hour with protein A-conjugated Dynabeads (Life Technologies), after which the beads were washed with lysis buffer at incremental KOAc concentrations (100 mM, 120 mM, 150 mM, and 60 mM NaOAc). Co-immunoprecipitated proteins were eluted with loading buffer and protein co-purification was visualized by western blot.

For the Chs2 alkaline phosphatase assay, metaphase arrested cells were collected and native protein extracts were prepared following the immunoprecipitation protocol. The protein extracts were incubated with alkaline phosphatase (Anthartic phosphatase, New England Biolabs) in EBX reaction buffer (50mM HEPES/KOH pH 7.5, 100mM KCl, 2.5mM MgCl_2_, 0.25% Triton X-100, 1M DTT with protease inhibitors (8 μg/mL (leupeptin, pepstatin, aprotinin)), 1 mM PMSF, 1.25mg/ml of benzamidin, 1 tablet of complete protease inhibitor without EDTA (Roche)) at 37°C for 15 minutes. To inhibit the alkaline phosphatase 2XPhosStop (Roche) was used.

To purify the PP2A-Cdc55 complex, a strain with the tandem affinity protein (TAP) epitope, TAP-HA-Cdc55 was used. Cells from 2 L cultures were arrested in metaphase by Cdc20 depletion, frozen in liquid nitrogen and stored at -80°C. Cells were resuspended in lysis buffer (0.1 M NaCl, 50 mM Tris-HCl pH 7.5, 1.5 mM MgCl_2_ and 0.15% NP-40, 8 μ mL of protease inhibitors (leupeptin, pepstatin, aprotinin), 1 mM PMSF, 1.25mg/ml of benzamidin, 1 tablet of complete protease inhibitor without EDTA (Roche)) and protein extracts prepared by mechanical lysis. Protein extracts were clarified by centrifugation at 20.000 g and incubated with 0.4 mL IgG-Sepharose beads for 1 hour. The beads were extensively washed and incubated with 3 μL of TEV protease at 16°C for 2 h to remove the CBP-HA-Cdc55 from the beads. The purified PP2A-Cdc55 was stored at -80°C and used for the phosphatase assays in Fig 3d and Fig. S5f.

The Chs2-HA and Esp1-HA were purified following the co-immunoprecipitation protocol as above but using Pierce HA-magnetic beads and eluted from the beads using 1mg/mL of HA peptide. For the phosphatase assays with purified PP2A-Cdc55, eluted Chs2 or Esp1 were incubated with phosphatase reaction buffer (50 mM Tris-HCl pH7.4, 0.1 mM EGTA, 1 mM β-mercaptoethanol, 1mg/mL BSA) and the purified CBP-HA-Cdc55 at 30°C for 20-40 minutes. Reactions were terminated by adding SDS-PAGE loading buffer. Proteins were separated by electrophoresis, transferred to PVDF membranes and phosphorylation levels detected with the anti-phospho Ser/Thr antibody (Cell Signaling) using the Supersignal West Atto Ultimate Sensitivity Substrate (Pierce). Anti-HA western blots were then performed to analyze and quantify the amount of Chs2, Esp1 and Cdc55. The phosphorylation levels were normalized against Cdc55 and Chs2 or Esp1. Proteins were quantified using Fiji software[74] and values of the mean and SEM calculated.

For radioactive ^32^P kinase assays, Clb2-Pk immunoprecipitations were performed as above and incubated with α-Pk clone SV5-Pk1 (AbSerotec) antibody. Beads were washed with ten volumes of lysis buffer and twice with the kinase reaction buffer (50 mM Tris-HCl, pH7.4, 10 mM MgCl_2_, 1 mM DTT). The kinase reaction (50 mM Tris-HCl, pH7.4, 10 mM MgCl_2_, 1 mM DTT, 5 mM β-glycerophosphate, 25 μM ATP, 10 mCi/mL ^32^gamma-ATP and 1 μg of Streptag-Chs2) was incubated at 30°C for 30 minutes. Kinase assays were stopped by placing the tubes on ice. The supernatant containing the phosphorylated substrate was separated from the magnetic beads and stored at -80°C. An aliquot of the kinase assay was mixed with SDS-PAGE loading buffer, proteins were separated by electrophoresis, transferred to nitrocellulose membranes and radioactivity detected in a Typhoon FLA950 (GE Healthcare). Immunopurified protein was quantified by western blot and the membrane was stained with Coomassie to detect the recombinant substrate. Proteins were quantified using Fiji software.

For *in vitro* phosphatase assays, cells containing HA_3_-*Cdc55-ED* and HA_3_-Cdc55 were collected from metaphase arrested cells. Immunoprecipitation was performed as above but without phosphatase inhibitors. Beads were washed twice with phosphatase buffer and incubated with the phosphatase reaction (50 mM Tris-HCl pH7.4, 0.1 mM EGTA, 1 mM β-mercaptoethanol, 1mg/mL BSA and the ^32^P-Streptag-Chs2) at 30°C for 40 minutes. Reactions were terminated by adding SDS-PAGE loading buffer. Proteins were separated by electrophoresis, transferred to nitrocellulose membranes and radioactivity was detected in a Typhoon FLA950 apparatus (GE Healthcare). Later, the membrane was used to analyze and quantify the amount of protein immunoprecipitated by western blot. Finally, the membrane was stained with Coomassie to detect the recombinant Chs2 substrate. Proteins levels were quantified using Fiji software and values of the mean and SEM calculated.

The *GAL1-YFP-Chs2* and *GAL1-YFP-Chs2-S133A* were purified following the co-immunoprecipitation protocol as above but using GFP-Trap beads (Chromotek). Proteins were separated by electrophoresis, transferred to PVDF membranes and phosphorylation levels detected with the anti-phospho Ser/Thr antibody (Cell Signaling) using the Supersignal West Atto Ultimate Sensitivity Substrate (Pierce). Anti-GFP western blots were then performed to analyze and quantify the amount of Chs2. The phosphorylation levels were normalized against total Chs2. Protein levels were quantified using Fiji software and values of the mean and SEM calculated.

### Microscopy techniques

Synchronized cells for time-lapse experiments were deposited in chambers (Nunc Lab-Tek) containing concanavalin A/PBS 1 mg/mL. Images were captured every 2 minutes. Different z-stacks at 0.7-μm intervals were taken and projected onto a single image per channel. A Zeiss-Apotome epifluorescence microscope with an HXP 120C fluorescent lamp and a Carl Zeiss Plan-Apochromat 63x N.A 1.40 oil immersion lens were used. The filters used were Cy3, GFP, and DAPI. For the GFP-Ras2 experiments the Apotome was disabled. For the Cyk3-GFP time-lapse experiments, a Carl Zeiss LSM880 confocal microscope, with a 63x N.A. objective was used. Images were acquired using ZEN software. For the time-lapses in figure 5, a Leica SP8 confocal microscope with a 63X objective was used. Images were acquired using LAS software and quantified and processed using Fiji software. To determine the symmetry of the IPCs proteins, the bud neck length was measured, the central point calculated and manually established whether the central point was inside (symmetry) or outside (asymmetry) the dot signal.

Calcofluor staining was performed with calcofluor white MR2 (Fluorescent brightener 28, Sigma) in living cells. Synchronized cells were incubated with 50 μg/mL calcofluor upon release from the metaphase arrest. Images were acquired, quantified and processed as for the time-lapse experiments.

Phalloidin staining was performed on fixed cells. Cells collected during the time-course experiments were pre-fixed with PBS containing 3.7% formaldehyde and 0.1% Triton X-100 at 25°C for 10 minutes. Cells were then washed with PBS1 and fixed with 3.7% formaldehyde at 25°C for 1 hour. After fixation, cells were washed twice with PBS1 and sedimented onto a multi-well slide previously incubated with poly-L-lysine. Cells were stained with a PBS solution containing 50 U/mL rhodamine phalloidin R415 (Life Technologies) for 2 hours. Cells were washed twice with PBS1. Mounting medium containing DAPI (Vectashield) was added, cells were covered with a coverslip and sealed with nail polish. Images were acquired, quantified and processed as for the time-lapse experiments.

For electron microscopy, cell samples were fixed with 0.2 M phosphate buffer (without salts) pH 7.4, containing 2.5% glutaraldehyde at 25°C, for 1 hour. Cells were washed three times with phosphate buffer without glutaraldehyde and rinsed with milli-Q water and post-fixed with 1% osmium tetroxide for 2 hours. They were then dehydrated in an acetone series (10%, 20%, 30%, 40%, 60%, 80%, and 100%) for 15-20 minutes. Ultrathin sections of 60-nm thickness were obtained using a UC6 ultramicrotome (Leica Microsystems, Austria) and stained with 2% uranyl acetate and lead citrate. Sections were observed in a Jeol EM J1010 apparatus (Jeol, Japan) and images were acquired at 80 kV with a 1k CCD Megaview camera.

### SILAC labeling and phosphoproteomic analysis

Y859 (*MATa lys2Δ::TRP1, arg4Δ::HIS3 MET-CDC20::LEU2*) and Y858 (as Y859, but *cdc55Δ* ) cells were grown in free-methionine minimum media containing ^13^C_6_-lysine and -arginine (heavy, Cambridge Isotope Laboratories Inc., US) or unmodified arginine and lysine (light), respectively. Protein extracts were prepared by mechanical lysis using glass beads in presence of protein inhibitors (Complete EDTA-free, Roche) and 2x phosphatase inhibitors PhosStop (Roche). Cell lysates were mixed 1:1 and digestion with trypsin was performed. The phosphopeptides were pre-enriched using TiO_2_ chromatography (GL Sciences Inc) followed by SIMAC purification. The mono-phosphorylated peptide fraction from the SIMAC enrichment was further subjected to a second TiO_2_ purification and pre-fractionated by HILIC chromatography (Hydrophilic Interaction Liquid Chromatography). The peptides were then analyzed by LC-MS/MS using a LTQ-Orbitrap Fusion Tribride mass spectrometer (Thermo Fisher Scientific). Peptide identification was performed using Proteome Discoverer v1.4.1.14 (Thermo Scientific) and search against Swiss Prot/Uniprot *Saccharomyces cerevisiae* database. Peptide quantification from SILAC labels was performed with Proteome Discoverer v1.4.

### Other techniques

Protein extracts for western blots were obtained by TCA protein extraction. For the phostag gels, 10 μM Phostag (Wako) 5-8% gels containing 10 mM MnCl_2_ were used. Protein gels were washed with 1 mM EDTA before transferring protein to remove the manganese. Proteins were quantified using Fiji software, and values of the mean and SEM calculated. To calculate the ratio of phosphorylated/dephosphorylated Chs2, the faster migration band corresponds to the dephosphorylated Chs2 (as shown in the alkaline phosphatase experiment Fig. S5e). The slower migration band observed in metaphase is considered the phosphorylated Chs2. For each triplicate, the signals were normalized against the higher signal in the blot. To assign the cells to metaphase, anaphase and cytokinesis we took in consideration the percentage of anaphase spindles and the FACS profiles (not shown). Time-points with intermediate percentages of anaphase spindles were not used in the quantification.

Antibodies used for western blots were α-HA clone 12CA5 (Roche), α-myc 9E10 (Babco), α-FLAG clone M2 (Sigma), α-Pk clone SV5-Pk1 (AbSerotec), α-Clb2 (y-180) sc-907 (Santa Cruz Biotechnology), α-tubulin clone YOL1/34 (Serotec), α-phosphoglycerate kinase (Life Technologies), α-HA rabbit (Sigma), α-Pgk1 (Invitrogen), and α -Chs2[17]. The secondary antibodies were: α-Mouse-HRP (GE Healthcare), α-Rabbit-HRP (GE Healthcare), and α-Goat-HRP (GE Healthcare). Antibodies used for immunofluorescence were: α-HA clone 12CA5 (Roche), α-Cdc14 (yE-17) sc-12045 (Santa Cruz Biotechnology), α-tubulin clone YOL1/34 (AbSerotec), and α-Cdc11 (Santa Cruz Biotechnology). The secondary antibodies were Cy3-labeled α-mouse (GE Healthcare), fluorescein-conjugated α-rat (Millipore), Cy3-labeled α-goat (GE Healthcare), red TEXAS α-rabbit (Jackson Laboratories) and 488 α-mouse (Life Technologies).

### Statistical analysis

All experiments were done at least three times. Statistical analyses were performed with Prism5. Student’s T-test was used to analyze the p-values of the different assays compared. A p-value <0.05 was considered statistically significant, p<0.0001 ****, p<0.001 ***, p < 0.01 **; p<0.05 *.

## Acknowledgements

We thank the optical microscopy unit of IDIBELL, the electron microscopy units and the proteomic services from IDIBELL and CCiT-UB for their support. The *cdc55-aid* mutant strain was a kind gift of Adam Rudner and the *GAL-CHS2-YFP* containing plasmids were a kind gift of Foong May Yeong. We thank all the members of our laboratory for discussing the work and for their critical reading of the manuscript. We thank CERCA Program/Generalitat de Catalunya for institutional support. EQ is funded by the grants BFU2016-77975-R (co-funded by the European Regional Development Fund, ERDF, a way to build Europe) from the Spanish Ministry of Economy, Industry and Competitiveness (MINECO) and PID2019-109027GB-I00 from the Spanish Ministry of Science, Innovation and Universities (MCIU). ASD was supported by the grant PID2019-106745GB-I00 from the Spanish Spanish Ministry of Science, Innovation and Universities (co-funded by the European Regional Development Fund) and a grant from the Consejería de Universidades, Investigación, Medio Ambiente y Política Social del Gobierno de Cantabria.

## Author contribution

YMR, DV, OVC, MF and EQ performed the experiments. YMR, ASD and EQ designed the experiments and interpreted the data. YMR and EQ wrote the manuscript. All authors read and discussed the manuscript.

## Supplemental Figure legends

**Figure S1. Septin structures are not altered in the absence of Cdc55.** Cycling cells from strains Y1588 and Y1589 were fixed and immunofluorescence *in situ* performed for septins Cdc11 and Shs1-HA visualization. α -Cdc11 and α-HA (12CA5) antibodies were used. Representative images of Cdc11 and Shs1 septin’ structures from *CDC55* and *cdc55Δ* cells are shown. Scale bar, 1 µm.

**Figure S2. Actin is polarized to the new bud site before cytokinesis completion in *cdc55Δ* cells.** Strains Y1434 and Y1435 were arrested in metaphase by Cdc20 depletion and released into anaphase by Cdc20 re-induction. Formaldehyde-fixed cells were stained with 50 U/mL of rhodamine-phalloidin. (a) Representative images of the actin staining and Myo1-GFP signals from *CDC55* and *cdc55Δ* at different cell cycle stages (metaphase, anaphase, cytokinesis and the next S phase) are shown. (b) Quantifications and representative images of the cells with re-polarized actin at the new bud site are shown. Scale bar, 1 μm.

**Figure S3. Myo1 asymmetry contraction in G1-synchronized cells in absence of Cdc55.** Strains Y1631 (N=15) and Y1761 (N=12) were arrested in G1 by α actor addition and released into cell cycle progression by pheromone removal. Cells were fixed with formaldehyde before taking images. Quantifications of the cells with asymmetric Myo1 contraction are represented (left panel). Representative images of Myo1 asymmetric contraction are shown (right panel). Scale bar, 1 μm.

**Figure S4. Proper cytoplasm separation during membrane abscission in the absence of Cdc55.** Strains Y1717 and Y1708 were arrested in metaphase by Cdc20 depletion and released into anaphase by Cdc20 re-induction. Time-lapse images were captured every 2 minutes. Membrane abscission was followed by visualization of the GFP_3_-Ras2 signal, and Myo1-tdTomato was used as a control for cytokinesis progression. Representative images from *CDC55* (N=12) and *cdc55Δ* (N=13) cells are shown (left panel). Images are taken from the best z-stack to visualize the division site. Scale bar, 1 μm. Quantification of the cumulative percentage of cells abscised is represented (right panel).

**Figure S5. PP2A-Cdc55 participates in the dephosphorylation of IPCs proteins during cytokinesis.** (a-d) Phosphorylation changes of Hof1, Inn1, Cyk3 and Iqg1 in absence of Cdc55. Strains Y1314, Y1394, Y1639, Y1640, Y1437, Y1438, Y1491 and Y1497 were arrested in metaphase and released into anaphase by Cdc20 depletion and re-addition. Protein phosphorylations were analyzed by western blot. Pgk1 and Myo1-Flag levels were used as a loading control. Mitosis progression was followed by analyzing the anaphase spindle elongation by *in situ* immunofluorescence. At least 100 cells were scored at each time-point. Quantifications of the western blots were performed using Fiji Software and means and SEMs are represented. Student’s unpaired t-test analyses were carried out using the Prism5 program. (e) Alkaline phosphatase assay for Chs2. Native protein extracts were prepared from Y1318 cells arrested in metaphase and incubated with alkaline phosphatase and alkaline phosphatase’s inhibitor as indicated. PhosStop was used as the alkaline phosphatase’s inhibitor. Chs2 phosphorylation was analyzed by western blot in Phos-tag gels. (f) Separase is not dephosphorylated by PP2A-Cdc55. Metaphase arrested cells of the strain Y695 was used to purify the PP2A-Cdc55 complex by TAP purification. Esp1-HA was purified from metaphase-arrested cells of strain Y1748. Purified Esp1-HA was incubated with the PP2A-Cdc55 complex at the indicated times and the separase phosphorylation levels were detected by western blot using the anti-phospho Ser/Thr antibody. Representative images of one phosphatase assay are shown. Protein levels were quantified using Fiji software. Quantifications of the remaining separase phosphorylation signal normalized to the amount of Cdc55 and Esp1 are shown. Means and SEMs of three phosphatase assays are represented. Student’s unpaired t-test analysis was carried out using the Prism5 program.

**Figure S6. PP2A-Cdc55 is required for proper IPCs localization and contraction at the division site.** Strains Y1572, Y1606, Y1454, Y1608, Y1306, Y1578, Y1574, Y1604, Y1576 and Y1575 were synchronized into anaphase by Cdc20 depletion and re-addition, and time-lapse images were captured every 2 minutes. Myo1-tdTomato was used as a control for cytokinesis progression. (a-e) Representative images of the indicated GFP-tagged IPCs proteins and Myo1 contractions from *CDC55* and *cdc55Δ* cells are shown. Iqg1 and Inn1 contraction at the bud neck are normal in the absence of Cdc55. By contrast, Hof1 and Cyk3 contractions are longer in the absence of Cdc55. Scale bar, 1 μm.

