## Supplementary figures and images for "PP2A-Cdc55 phosphatase coordinates actomyosin ring contraction and septum formation during cytokinesis"

### FigureS1

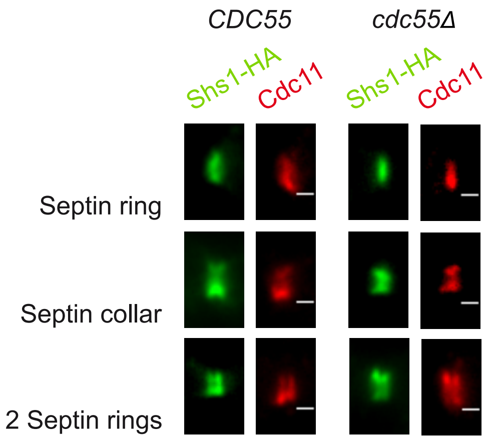

### FigureS2

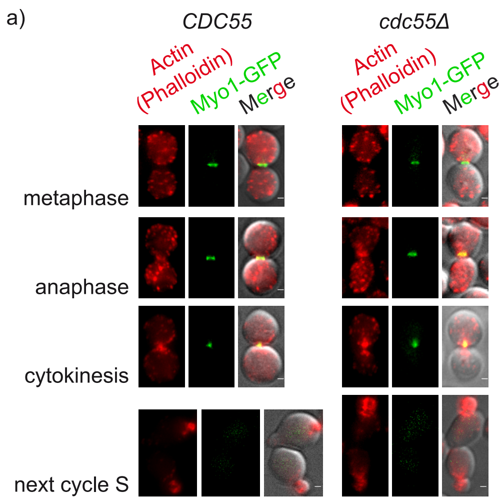

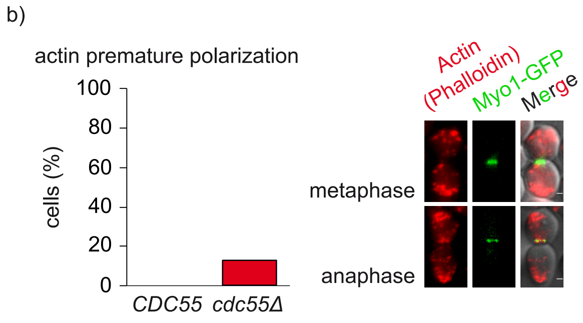

### FigureS3

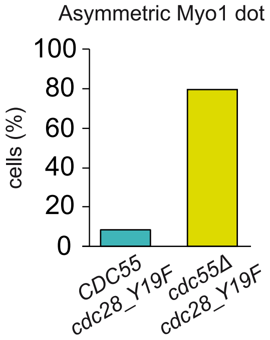

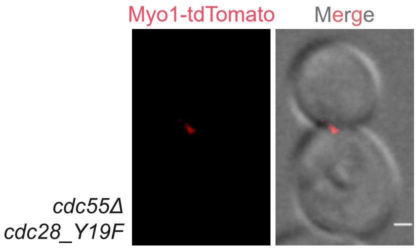

### FigureS4

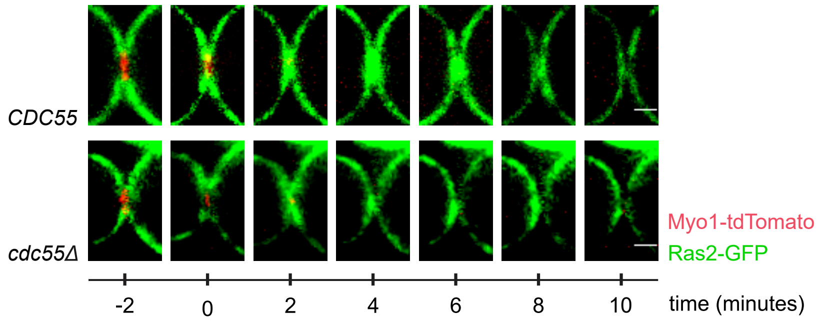

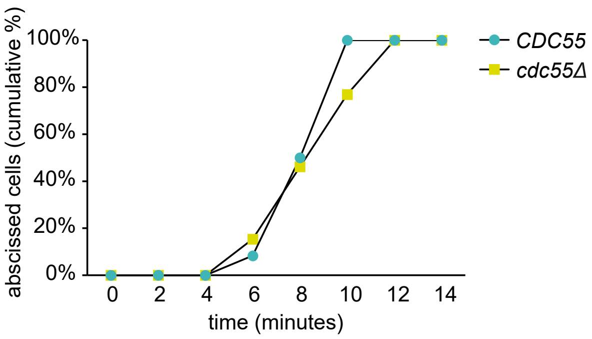

### FigureS5

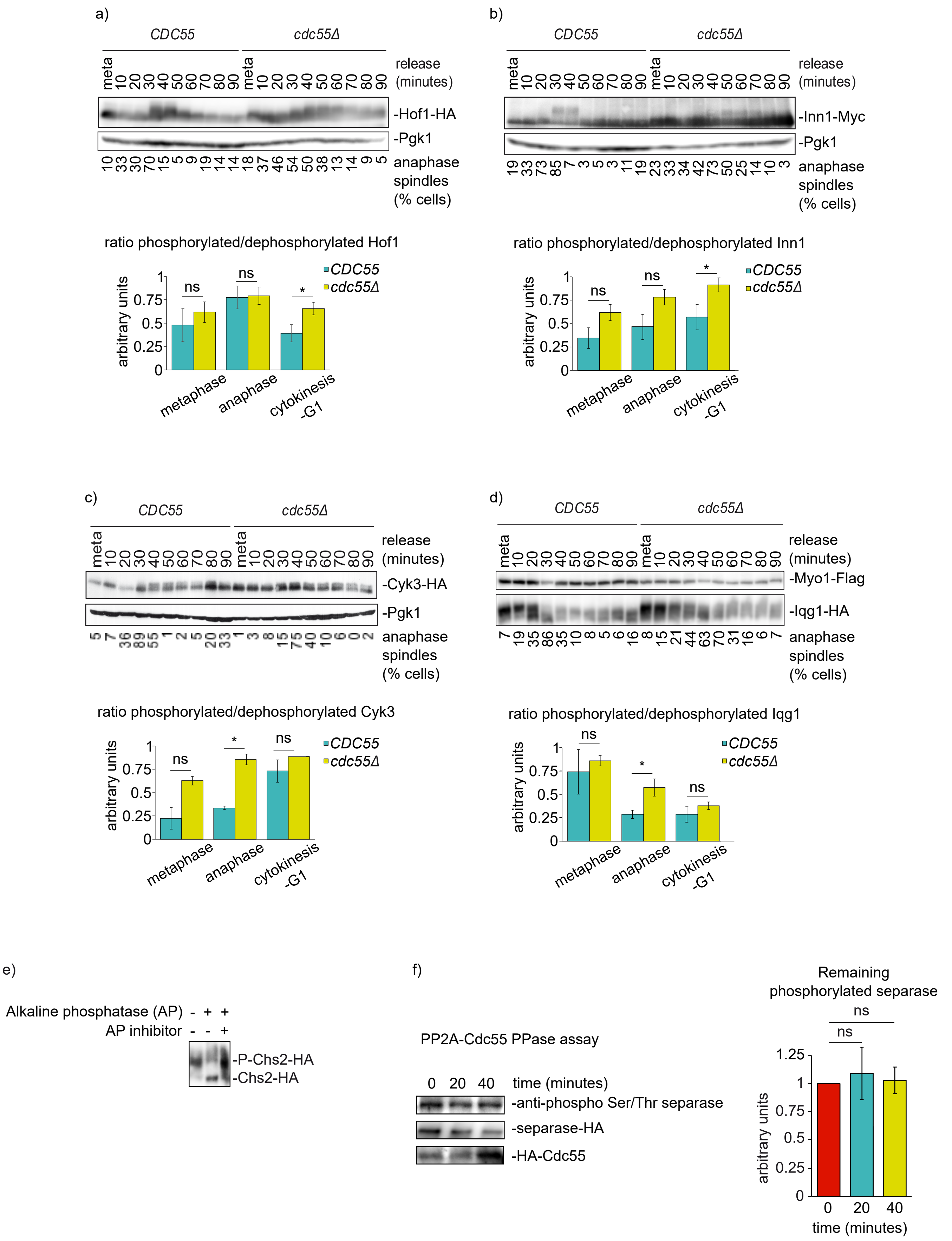

### FigureS6

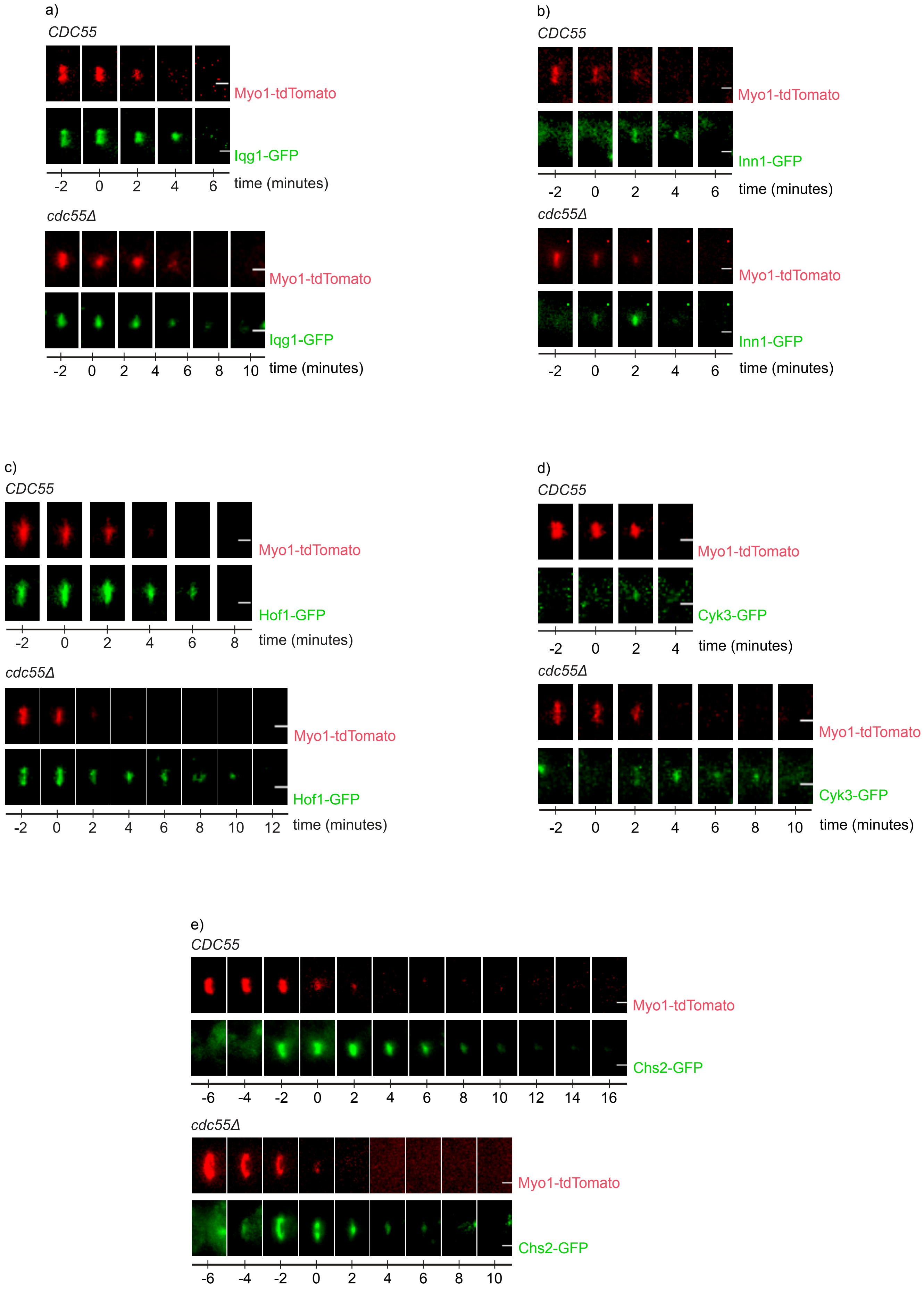
