## Supplementary material for "PP2A-Cdc55 phosphatase coordinates actomyosin ring contraction and septum formation during cytokinesis": TableS1

**Table S1. List of strains used in this study**

| Strain | Genotype | Origin |
|---|---|---|
| W303 | *MATa ade2-1 trp1-1 can1-100 leu2-3,112 his3-11,15 ura3 GAL psi+* | Matt Sullivan |
| Y695 | *MATa CDC14-Pk9 TAP-HA-CDC55 GAL1-CDC20* | This study |
| Y824 | *MATα MET-HA3-CDC20* | This laboratory |
| Y844 | *MATα cdc55Δ* | This laboratory |
| Y1306 | *MATα MET-HA3-CDC20 MYO1-tdTOMATO SPC42-GFP HOF1-GFP* | This study |
| Y1314 | *MATa MET-HA3-CDC20 HOF1-HA6* | This study |
| Y1315 | *MATα MET-HA3-CDC20 CHS2-HA6 cdc55Δ* | This study |
| Y1318 | *MATa MET-HA3-CDC20 CHS2-HA6* | This study |
| Y1394 | *MATα MET-HA3-CDC20 HOF1-HA6 cdc55Δ* | This study |
| Y1434 | *MATa MET-HA3-CDC20 MYO1-GFP cdc55Δ* | This study |
| Y1435 | *MATa MET-HA3-CDC20 MYO1-GFP* | This study |
| Y1437 | *MATa MET-HA3-CDC20 CYK3-HA6* | This study |
| Y1438 | *MATa MET-HA3-CDC20 CYK3-HA6 cdc55Δ* | This study |
| Y1454 | *MATa MET-HA3-CDC20 MYO1-tdTOMATO SPC42-GFP INN1-GFP* | This study |
| Y1491 | *MATa MET-HA3-CDC20 MYO1-FLAG IQG1-HA6 HOF1-MYC9* | This study |
| Y1497 | *MATα MET-HA3-CDC20 MYO1-FLAG IQG1-HA6 cdc55Δ* | This study |
| Y1512 | *MATa MET-HA3-CDC20 MYO1-tdTOMATO SPC42-GFP cdc55Δ* | This study |
| Y1516 | *MATa MET-$HA_3$-CDC20 CHS2-GFP SPC42-GFP* | This study |
| Y1567 | *MATa MET-HA3-CDC20 CHS2-HA6 PK3-CDC55* | This study |
| Y1572 | *MATa MET-HA3-CDC20 MYO1-tdTOMATO SPC42-GFP yE-GFP-IQG1* | This study |
| Y1574 | *MATa MET-HA3-CDC20 MYO1-tdTOMATO SPC42-GFP CYK3-GFP* | This study |
| Y1575 | *MATa MET-HA3-CDC20 MYO1-tdTOMATO CHS2-GFP cdc55Δ* | This study |
| Y1576 | *MATα MET-HA3-CDC20 MYO1-tdTOMATO SPC42-GFP CHS2-GFP* | This study |
| Y1578 | *MATa MET-HA3-CDC20 MYO1-tdTOMATO HOF1-GFP cdc55Δ* | This study |
| Y1588 | *MATa MET-HA3-CDC20 SHS1-HA6* | This study |
| Y1589 | *MATa MET-HA3-CDC20 SHS1-HA6 cdc55Δ* | This study |
| Y1596 | *MATα MET-HA3-CDC20 MYO1-tdTOMATO chs3Δ cdc55Δ* | This study |
| Y1604 | *MATa MET-HA3-CDC20 MYO1-tdTOMATO SPC42-GFP CYK3-GFP cdc55Δ* | This study |
| Y1605 | *MATα MET-HA3-CDC20 MYO1-tdTOMATO chs3Δ* | This study |
| Y1606 | *MATa MET-HA3-CDC20 MYO1-tdTOMATO SPC42-GFP yE-GFP-IQG1 cdc55Δ* | This study |
| Y1608 | *MATa MET-HA3-CDC20 MYO1-tdTOMATO SPC42-GFP INN1-GFP cdc55Δ* | This study |
| Y1631 | *MATa MYO1-tdTOMATO CDC15-eGFP cdc55Δ cdc28-Y19F* | This study |
| Y1639 | *MATa MET-HA3-CDC20 CYK3-HA6 INN1-Myc9* | This study |
| Y1640 | *MATa MET-HA3-CDC20 CYK3-HA6 INN1-Myc9 cdc55Δ* | This study |

| | | |
|---|---|---|
| Y1652 | *MATa MET-HA3-CDC20 MYO1-tdTOMATO cdc55Δ URA3::HA3-CDC55* | This laboratory |
| Y1653 | *MATa MET-HA3-CDC20 MYO1-tdTOMATO cdc55Δ URA3::HA3-CDC55-T174E_S301D* | This laboratory |
| Y1708 | *MATa MET-HA3-CDC20 MYO1-tdTOMATO 3GFP-RAS2 cdc55Δ* | This study |
| Y1717 | *MATa MET-HA3-CDC20 MYO1-tdTOMATO 3GFP-RAS2* | This study |
| Y1728 | *MATa ADH1-AtTIR1-MYC9 hof1-aid* | Sánchez-Díaz A laboratory |
| Y1730 | *MATa GAL1-UBR1 ADH1-AtTIR1-MYC9 td-cyk3-aid* | Sánchez-Díaz A laboratory |
| Y1747 | *MATa GAL1-UBR1 ADH1-AtTIR1-MYC9 td-cyk3-aid cdc55Δ* | This study |
| Y1748 | | |
| Y1749 | *MATa ADH1-AtTIR1-MYC9 hof1-aid cdc55Δ* | This study |
| Y1761 | *MATa MYO1-tdTOMATO CDC15-eGFP cdc28-Y19F* | This study |
| Y1788 | *MATα MET-HA3-CDC20 MYO1-tdTOMATO SPC42-GFP pGPD1-OsTIR1-MYC9 cdc55-aid* | This study |
| Y1893 | *MATa MET-HA3-CDC20 MYO1-tdTOMATO SPC42-GFP GAL-CHS2-S133E-YFP* | This study |
| Y1894 | *MATa MET-HA3-CDC20 MYO1-tdTOMATO SPC42-GFP GAL-CHS2-S133E-YFP cdc55Δ* | This study |
| Y1895 | *MATa MET-HA3-CDC20 MYO1-tdTOMATO SPC42-GFP GAL-CHS2-YFP* | This study |
| Y1896 | *MATa MET-HA3-CDC20 MYO1-tdTOMATO SPC42-GFP GAL-CHS2-YFP cdc55Δ* | This study |
| Y1898 | *MATa MET-HA3-CDC20 MYO1-tdTOMATO SPC42-GFP GAL-CHS2-S133A-YFP cdc55Δ* | This study |
| Y1900 | *MATa MET-HA3-CDC20 MYO1-tdTOMATO GAL-CHS2-S133A-YFP* | This study |
| Y1901 | *MATa MET-HA3-CDC20 MYO1-tdTOMATO GAL-CHS2-YFP* | This study |
| Y1909 | *MATa MET-HA3-CDC20 HOF1-mCherry SPC42-GFP GAL-CHS2-YFP* | This study |
| Y1911 | *MATa MET-HA3-CDC20 HOF1-mCherry SPC42-GFP GAL-CHS2-S133E-YFP* | This study |
| Y1912 | *MATa MET-HA3-CDC20 HOF1-mCherry SPC42-GFP GAL-CHS2-YFP cdc55Δ* | This study |
| Y1913 | *MATa MET-HA3-CDC20 HOF1-mCherry SPC42-GFP GAL-CHS2-S133A-YFP cdc55Δ* | This study |
